# Quantification of histone H1 subtypes using targeted proteomics

**DOI:** 10.1101/2024.01.20.576464

**Authors:** Jordi López-Gómez, Laura Villarreal, Marta Andrés, Inma Ponte, Blanca Xicoy, Lurdes Zamora, Marta Vilaseca, Alicia Roque

## Abstract

Histone H1 is involved in the regulation of chromatin structure. Human somatic cells express up to seven subtypes. The variability in the proportions of somatic H1s (H1 complement) is one evidence supporting their functional specificity. Alterations in the protein levels of different H1 subtypes have been observed in cancer, suggesting their potential as biomarkers and that they might play a role in disease development. We have developed a mass spectrometry based (MS) parallel reaction monitoring (PRM) assay suitable for the quantification of H1 subtypes. Our PRM method is based on the quantification of unique peptides for each subtype, providing high specificity. Evaluation of the PRM performance on three human cell lines showed high reproducibility and sensitivity. Quantification values agreed with the electrophoretic and Western blot data, indicating the accuracy of the method. We used PRM to quantify the H1 complement in peripheral blood samples of healthy individuals and chronic myeloid leukemia (CML) patients. Our preliminary data revealed differences in the H1 complement between responders and non-responder CML patients and suggest that the H1 content could help predicting imatinib response.

## Introduction

Histone H1 is a protein family involved in the regulation of chromatin structure and gene expression. In humans, it is composed of 11 subtypes or variants. Seven subtypes, H1.0-H1.5 and H1X, are expressed in somatic cells, while the remaining subtypes are germline-specific. Somatic subtypes are divided into two groups. H1.1-H1.5 belongs to the replication-dependent (RD) subtypes, whose transcription is associated with the histone locus body. H1.0 and H1X have different expression patterns during the cell cycle, which are known as replication-independent (RI) subtypes [1].

Histone H1 is composed of three structural domains: a short N-terminal, a conserved globular domain, and a long C-terminal. Both terminal domains are intrinsically disordered with a high number of basic residues [2]. RD subtypes have high sequence identity (65-86%), with H1.2-H1.4 being the more similar subtypes. In contrast, H1.1 has a lower sequence identity than the other RD subtypes [3]. Both RI subtypes have a more divergent amino acid sequence while conserving the enrichment in basic residues.

Histone H1 complement is defined as the proportions of the different H1 subtypes in a cell at a given moment. It is variable depending on the cell type, the cell cycle phase, and the time of development [4]. However, the regulatory mechanisms behind the differential expression of H1 subtypes are unknown. The H1 complement in human cell lines has been studied after H1 separation by capillary zone electrophoresis [5]. In mouse embryonic and adult tissues, the H1 complement has been determined after separating individual subtypes by chromatography (reviewed in [4]). In the latter case, separation was not always complete, and some reported values included more than one subtype.

Alterations in the H1 complement have been extensively described in cancer, suggesting its potential as a biomarker for this disease (Table 1 [6–20]). However, the analysis of the H1 complement composition is particularly challenging for several reasons: i) RD-subtypes have over 65% of sequence identity; ii) denaturing polyacrylamide gel electrophoresis (SDS-PAGE) of perchloric extracts does not resolve individual variants, as they have similar net charge and molecular weight; iii) Western blot and immunohistochemistry detection depends on antibody performance and are not suitable for multiplexing; iv) untargeted proteomic analysis should take into account unique specific peptides not shared between subtypes and would be intended for relative quantification of the protein subtypes between samples. There are untargeted proteomic approaches considering emPAI index number (exponentially modified abundance index[21]) or iBAQ (intensity based absolute-quantitation [22]), which could estimate the amount of protein, or in this case, histone subtypes in a sample. However strictly speaking, neither antibody-based techniques nor the above mentioned untargeted proteomics methods could accurately quantify the absolute amounts of different subtypes in the same sample. Therefore, there is an unmet need to develop an assay suitable for the unambiguous absolute quantification of individual subtypes that allows the determination of the H1 complement composition in cell lines and biological samples, which often contain limited protein amounts. PRM (Parallel reaction monitoring) is a MS-based proteomics technique which combined with the introduction of isotopically labeled selected peptide standards in known concentrations enables the quantification of low amounts of proteins with high accuracy, sensibility and reproducibility [23,24]. We therefore envisaged this technique for the absolute quantification of all somatic subtypes without ambiguity.

**Table 1.**
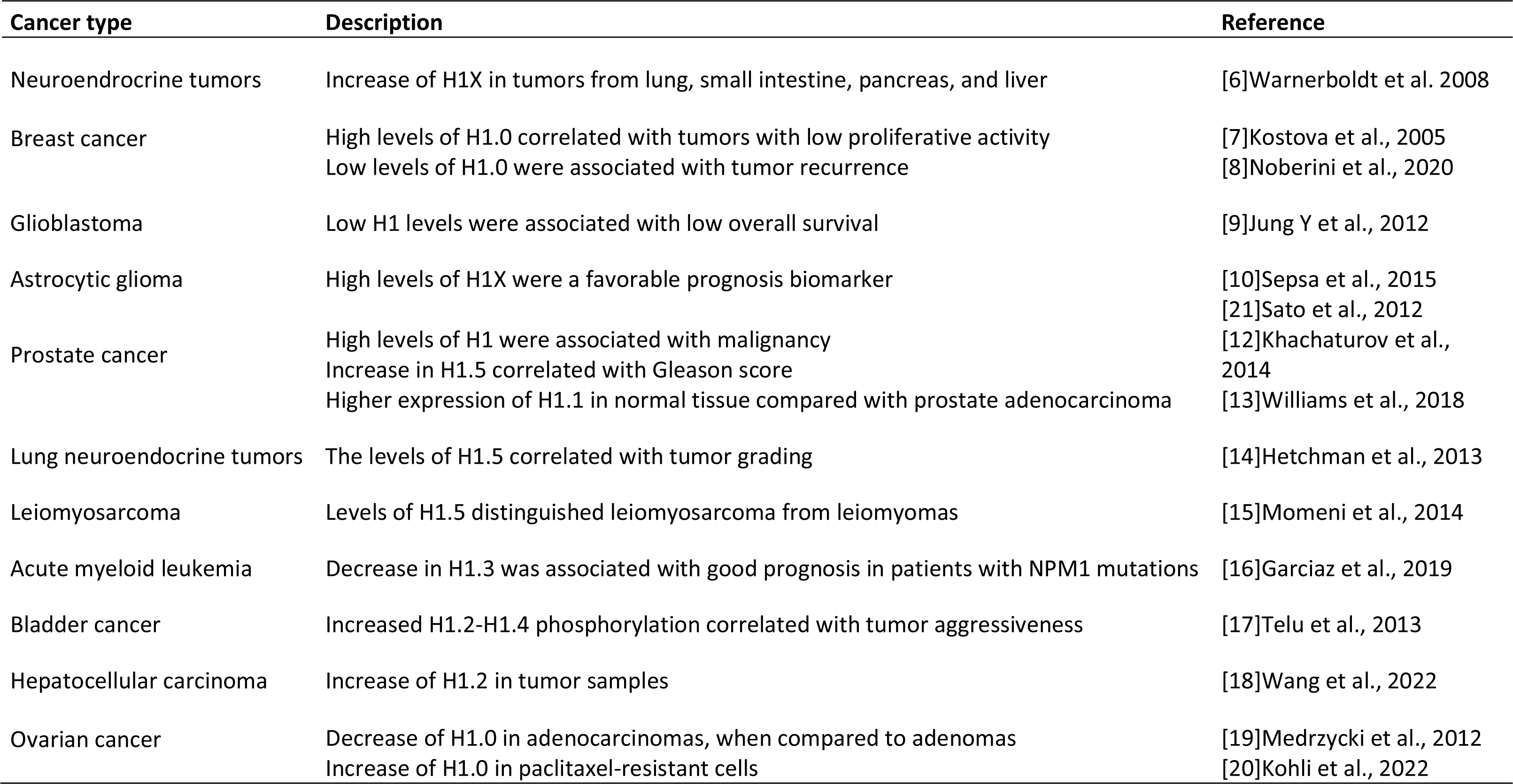
Alterations of the protein levels of H1 subtypes in tumor samples.

In this work, we report the design and standardization of a parallel reaction monitoring (PRM) assay, able to unambiguously quantify the seven H1 somatic subtypes from different sample types. We used this assay to characterize the histone H1 complement of three cancer cell lines and peripheral blood samples from healthy subjects.

As a proof of concept, we analyzed the H1 levels in a small cohort of chronic myeloid leukemia (CML) patients. CML is a hematological malignancy characterized by the reciprocal translocation between chromosomes 9 and 22 that form the Philadelphia chromosome. This alteration produces the BCR-ABL fusion protein, which is considered the driver mutation. Tyrosine kinase inhibitors (TKIs) are the standard treatment of CML and imatinib is the most used as first-line treatment.

However, approximately 20% of the patients are resistant to this therapy [25,26]. We found that the amount of total H1 was higher in patients that didn’t respond to imatinib than in patients with complete response to this drug. Non-responder patients had a characteristic H1 profile with low levels of H1.0, H1.1, and H1X.

## Materials and methods

### Synthetic peptides

Isotopically labeled synthetic peptides and light peptides corresponding to H1 somatic with purity > 95% were purchased from Synpeptide Co. Ltd (Shanghai, China) (Table S1). All peptides were supplied after HPLC purification and mass spectrometry verification.

### Cell culture

All the cell lines were grown at 37°C and 5% CO_2_ in their specific culture media supplemented with 10% fetal calf serum (Gibco) and 1% penicillin-streptomycin (Ddbiolab). Cervical carcinoma cells (HeLa) were cultured in DMEM Glutamax (Corning). For human breast cancer cells (T47D) and chronic myeloid leukemia cells (K562), the culture media was RPMI (Corning), supplemented with 2 mM L-glutamine (Ddbiolab).

### Peripheral blood samples

Human samples of peripheral blood from three healthy individuals and eight patients of CML were obtained following the ethical guidelines of the Germans Trias i Pujol Hospital. Blood samples of 10 mL were stabilized with EDTA and centrifuged at 800g for 10 minutes. The supernatant was recovered and centrifuged at 800g for 10 minutes. The cellular pellet was resuspended in 2.5-3 volumes of TLE buffer (NH_4_Cl 144 mM, NH_4_HCO_3_ 10 mM) and incubated in a rotating wheel for 10 minutes at room temperature to lyse the erythrocytes. The sample was centrifuged at 800g for 10 minutes, and the pellet corresponding to the white blood cells was resuspended in 0.5-2mL of TLE buffer. Finally, the cells were recovered by centrifugation at 800g for 10 min, and the dry pellet was stored at –80°C until further use.

### *In silico* analysis of peptide candidates

Protein sequences of human H1 somatic subtypes (Uniprot accession numbers: P07305, Q02539, P16403, P16402, P10412, P16401, Q92522) were used for simulation of proteolytic digestion with trypsin and Glu-C in the PeptideMass tool from Expasy (https://web.expasy.org/peptide_mass/). Unique peptides with less than 25 residues, and a monoisotopic mass of at least 600 Da were selected. The empirical suitability score (ESS) of the preselected peptides was extracted from the Peptide Atlas database (https://peptideatlas.org/) using the Human build (2023-01).

### Preparation of protein extracts

Cells were harvested by trypsinization and washed twice with phosphate-buffered saline (PBS) supplemented with a protease inhibitor cocktail (PIC, Roche). For obtaining linker histones, extraction with 5% perchloric acid was performed as previously described [3]. For preparing total histones, sulfuric extraction was performed using 0.2 M sulfuric acid instead of perchloric acid, using the same procedure described above.

For preparing total protein extracts, cells were resuspended in cold PBS, 0.1 mM DTT supplemented with PIC, and lysed by sonication in a Branson sonifier SFX250, at 40% power using three pulses of 30 seconds, alternating with 30 seconds on ice. Sodium chloride was added up to 500 mM and incubated in a rotating wheel for one hour. Afterward, the sample was centrifuged, and the supernatant dialyzed against the resuspension buffer in a mini tube-O-dialyzer (G-Biosciences) with a cut-off of 1 kDa. The supernatant contained a mixture of cellular proteins that were precipitated in 20% trichloroacetic acid overnight at 4°C. The sample was centrifuged at 16000 g for 15 minutes at 4°C, and the pellet was washed with cold acetone twice.

All the protein extracts were resuspended in 100 mM ammonium bicarbonate in a Lobind microcentrifuge tube (Eppendorf), dried by lyophilization, and stored at –20°C. Protein concentration was determined with BCA assay (Thermofisher).

### Western blot

Equivalent amounts of total histones (5 μg) or total protein extracts (20 µg) of the three cell lines were separated in a 12% denaturing polyacrylamide gel electrophoresis (SDS-PAGE) and electrotransferred to a polyvinylidene difluoride membrane (PVDF) (EMD Millipore) at 100V for 1h. Immunoblot analyses were performed with the conditions recommended by the manufacturer for the primary and secondary antibodies (Table S2). The specificity of primary antibodies has been validated using knock-down cell lines [27]. Blots were visualized with Clarity Western ECL substrate (Bio-Rad) in a Chemidoc imaging system (Bio-Rad). Histone H3 and tubulin were used as loading controls for total histone and total protein extracts, respectively.

### Sample preparation for proteomic analyses

Sample digestion method was optimized to minimize missed cleavages. For denaturation step, 8M Urea vs. heating at 95°C for 5min or 45min at 56°C with dithiothreitol (DTT) 2 mM with posterior carbamidomethylation were tested. For digestion three different Trypsin enzymes (Trypsin-Promega; Trypsin/LysC-Promega and fastTrypsin-Promega), 3 different time points incubation (24, 48 and 72h) and two enzyme: protein ratios (1:20 and 1:50) were tested.

For the final method, dry extracts were reconstituted in 50 mM NH_4_HCO_3_, pH 8.0, at a final concentration of 1 mg/mL. Samples were then divided in two to obtain tryptic and Glu-C peptides and diluted to final concentration of 0.1 mg/mL with 50 mM NH_4_HCO_3_ for Trypsin digestion and with PBS for Glu-C (Promega) digestion. DTT was added to a final concentration of 2 mM and incubated at 56°C for 45 min, after lowering the temperature of the samples to RT, iodoacetamide (IAM) was added to a final concentration of 5 mM with incubation for 30 min at RT in the dark.

DTT was added again to a final concentration of 2 mM to consume any unreacted IAM. The enzyme solution was added to a ratio of 1:20 (w/w, enzyme:protein) and left to react for 20 h at 37 °C, next day more enzyme was added, this time at a ratio 1:40 and left to react for 4 h at 37 °C. The digestion was stopped by adding trifluoroacetic acid to a final concentration of 1 %. Then samples were cleaned up with polyLC C18 tips and dried in the Speed-Vac. Peptides were reconstituted in 3% acetonitrile (ACN) and 1% formic acid (FA) to a final concentration of 1 mg/mL and then diluted 1:5 for MS analysis.

### Internal and external curve calibration preparation

For the calibration curve of the heavy peptides, the points of the curve were prepared adding different volumes of the stock of mixing heavy peptides and 3% ACN and 1% FA to a final volume of 20 μL in HPLC vials (Table S3). These curves were injected in triplicates using the nanoLC-MS/MS method run in the Orbitrap Eclipse described below; the injected volume was 6 μL. The external light peptides curves were prepared as follows: different volumes of light stock peptides were loaded in the evotips and diluted with the respective volume of 3% ACN and 1% FA to a final volume of 20 μL, before loading into the instrument (Table S4). The sample optimization and initial setup of the MS method to assess the linearity of each peptide, the limits of detection and quantification were performed using a Dionex-Orbitrap Lumos system, as described below (Table S5).

### nanoLC-MS/MS

nanoLC-MS/MS experiments were acquired in a two-event experiment, data dependent acquisition (Exp. 1) and targeted (Exp. 2), DDA-tMS, using both an Orbitrap Eclipse and an Orbitrap Fusion Lumos (Thermo Scientific) hyphenated to the EvosepOne nanoLC system (EVOSEP, Odense, Denmark) or to an Ultimate Dionex nanoLC System (Ultimate 3000, ThermoScientific).

The sample optimization part was driven in the Dionex-Orbitrap Lumos system, with the following specifications. The sample was loaded to 100 μm × 2 cm Acclaim PepMap100, 5 μm, 100 Å, C18 (Thermo Scientific) at a flow rate of 15 μL/min using a Dionex chromatographic system. Peptides were separated using a C18 analytical column (NanoEase MZ HSS T3 column, 75 μm × 250 mm, 1.8 μm, 100 Å, Waters) with 250 nl/min flow and 90 min run, comprising three consecutive steps with linear gradients from 1 to 35% B in 60 min, from 35 to 50% B in 5 min, and from 50 % to 85 % B in 2 min. The column outlet was directly connected to an Advion TriVersa NanoMate (Advion) fitted on an Orbitrap Lumos. The mass spectrometer was operated in DDA-tMS mode. For DDA experiment MS1 scans were acquired in the orbitrap with the resolution (defined at 200 m/z) set to 120,000, and the scan range was set to m/z 350-1200, with lock mass on (445.12 m/z). The top speed (most intense) ions per scan were fragmented by CID, with 35% of collision energy, and detected in the Orbitrap with 30k resolution. The ion count target value was 400,000 for the survey scan and 50,000 for the MS/MS scan. Target ions already selected for MS/MS were dynamically excluded for 30 seconds. For tMS experiment, selected ions were fragmented by HCD with 28% of collision energy and detected in the Orbitrap with 30k resolution. The maximum injection time was 200 ms and the AGC targeted 100,000. The spray voltage in the NanoMate source was set to 1.70 kV.

Final measurements were acquired using an EvosepOne coupled to an Orbitrap Eclipse mass spectrometer via an Easy-spray source interface and a stainless-steel emitter (EV-1086 EVOSEP). Total protein digests samples with tryptic or Glu-C heavy peptide stock were loaded onto the EVOTIP (EV2013) precolumn following the manufacturer’s instructions. We used a 15cm Evosep column (EV-1137, 150µm ID, 1.5 µm beads) installed on a Butterfly heater (Phoenix S&T) at 40°C. The Evosep One method was 15SPD Evosep (88 min gradient), which uses a 220 nl/min flow rate[28]. Mobile phase A was 0.1% formic acid (FA) in water, and mobile phase B was 0.1% formic acid (FA) in acetonitrile (ACN). The mass spectrometer was operated in MS1-tMS mode. MS1 scans were acquired in Orbitrap with 60K resolution, a scan range of 300-1200, with the AGC targeted 100,000 and maximum injection time of 50ms. For the tMS experiment, the selected ions were fragmented by HCD with 28% of collision energy and detected in the orbitrap with 30K resolution. Orbitrap Eclipse & Lumos Tune Application 3.5.3890 and 4.0.4091 and Xcalibur versions 4.5.445.18 and 4.6.67.17 were used to operate the instruments and to acquire data, respectively. In some peripheral blood samples, miscleavaged peptides were detected, so external calibration curves using light peptides were used to correct the quantification values. External calibration curves were not acquired the same day as the samples.

### Database searches for MS method optimization

A database search was performed with Proteome Discoverer software v2.3 (Thermo) using Sequest HT search engine and SwissProt database [Human release 2019 05 with contaminants database]. Searches were run against targeted and decoy databases to determine the false discovery rate (FDR). Search parameters included enzyme specificity, allowing for two missed cleavage sites, methionine oxidation, and N-terminus acetylation as dynamic modifications and carbamidomethyl in Cys as fixed modification. Peptide mass tolerance was 10 ppm and the MS/MS tolerance was 0.02 Da. Peptides with FDR < 1% were considered positive identifications with a high confidence level.

### Database searches and data processing for absolute protein quantification

We performed a database search with MaxQuant v1.6.14.0 (MQ) software with Andromeda as a search engine to build a spectral library in Skyline software. The database used in the search was a Fasta created with the 7 Histone H1 (H1.0, H1.1, H1.2, H1.3, H1.4, H1.5 and H1X). We ran a search against targeted and decoy databases to determine the false discovery rate (FDR). Search parameters included trypsin or GluC enzyme specificity, allowing for two missed cleavage sites, oxidation in M, acetylation in protein N-terminus, Ile (+7), Leu (+7), Lys (+8), and Ala (+4) as dynamic modifications and Carbamidomethyl in Cys as static modification. Peptide and MS/MS mass tolerance was 20 ppm. Quantitative targeted MS/MS analysis was performed using Skyline v20.2.0.343, an open-source software project [29]. A spectral library was generated in Skyline from database searches of the targeted MS/MS raw files with MaxQuant. We introduced the targeted peptides, with oxidation in Met and the heavy labels in Lys (+8), Leu (+7), Ile (+7), and Ala (+4). The final selected peptides were manually imported within Skyline. Peaks were picked in an automated fashion using the default Skyline peak picking model, with Savitzky–Golay smoothing. Peak areas integration was based on extracted ion chromatograms (XICs) of MS/MS fragment ions masses, typically y– and b-ions, matching to specific peptides present in the spectral library. All transitions and peak area assignments were manually validated. The ratio of Light/Heavy was obtained from the peak area integration from light and heavy peptides. Skyline output tables contain calibration curve data: replicate and ratio Light/Heavy (LH) for each peptide. We calculated the final histone subtype amount (ng) per total protein (µg). We used R statistical software to do all the calculations [R Core Team. (2014). R: A Language and Environment for Statistical Computing. Available online at: http://www.Rproject.org]. We first obtained the calibration curves for each LH (ratio Light/Heavy) for each peptide using lm function in R and obtained the slope and intercept. We used one point of the calibration curve for sample quantification. In human samples, quantitative values of H1 subtypes were expressed as ng H1 per milliliter of peripheral blood.

The mass spectrometry proteomics data have been deposited to the ProteomeXchange Consortium via the PRIDE[30] partner repository with the dataset identifier PXD050243.

## Data analysis

To evaluate the variations in histone percentages between PRM and SDS methods, we initially converted the data into logit form and fitted a linear model. This model incorporated a combination of the method, cell type, and histone group as explanatory variables. Then, we conducted a comparative analysis using the ‘glht’ function from the ‘multcomp’ package in R (http://multcomp.r-forge.r-project.org/). Finally, we accounted for multiple comparisons by applying the Benjamini-Hochberg correction. The receiver operating curve (ROC) and its derived parameters, Youden’s index, specificity, sensitivity, and AUC, were obtained using Epitools (https://epitools.ausvet.com.au/roccurves) and SRplot https://www.bioinformatics.com.cn/en.

## Results

### Peptide selection and PRM setup

MS-based Parallel Reaction Monitoring relies on the selection of unique peptides called proteotypic peptides, which meet specific criteria [31]. Absolute quantification is performed by adding to the sample the isotopically labeled proteotypic (SIS) peptides in known concentrations as internal standards. Proteotypic peptides are targeted by their m/z ratio in the mass spectrometer analyzer and subsequently fragmented, producing product ions. Quantification was performed using the sum of the area of the extracted ion chromatogram peak (XIC) from each product ion of the precursor peptide ions, referred to that of the corresponding SIS-peptide (Figure 1A)[23,24].

**Figure 1.**
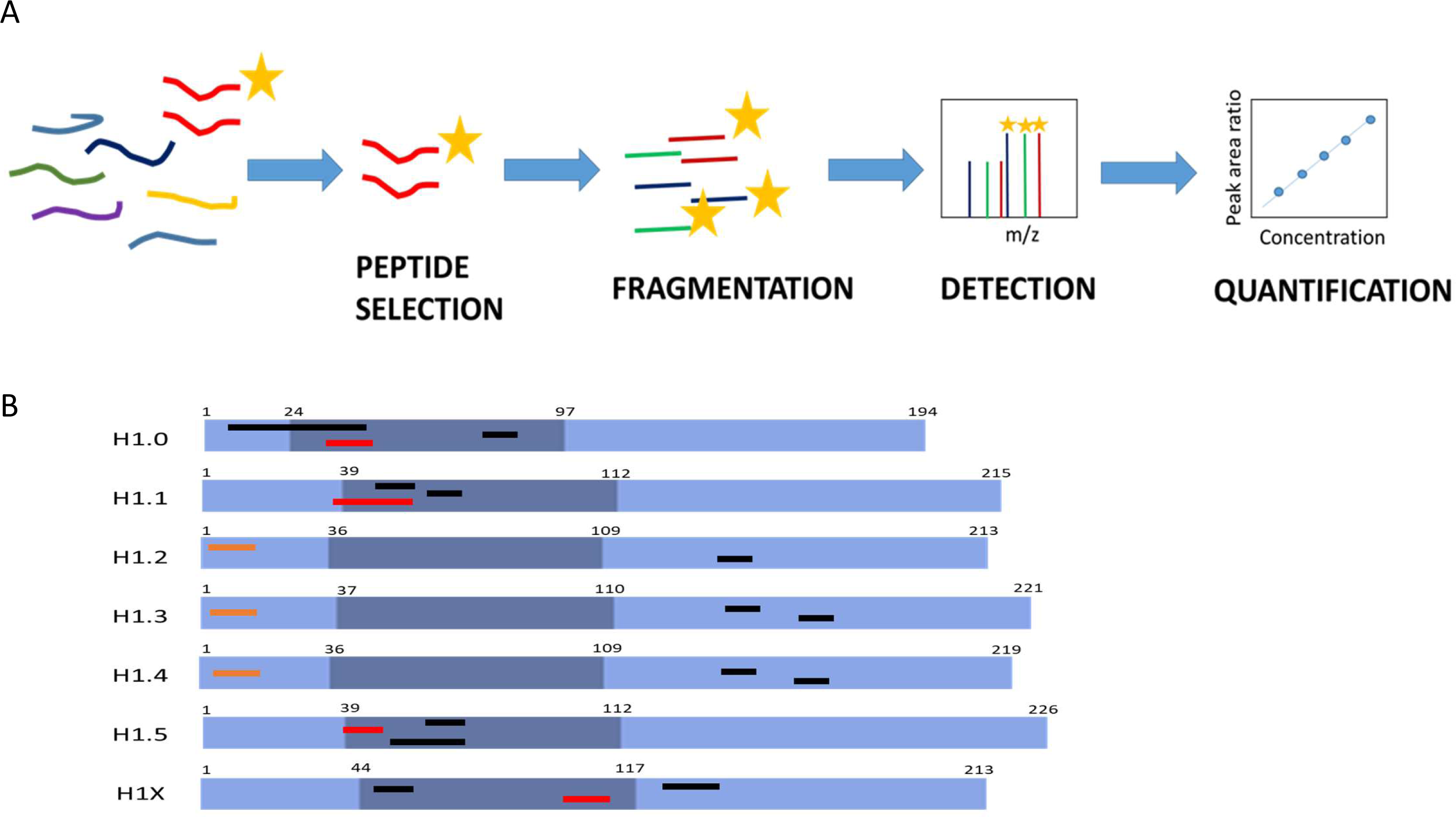
Parallel reaction monitoring assay setup. A. Scheme describing the procedure of a PRM assay. B. Representation of H1 somatic subtypes indicating the localization of the heavy peptides. Light blue regions correspond to the N– and C-terminal domains, while darker blue denotes the globular domain. The number corresponds to the residues at the border of each domain. Highlighted in black are the pre-selected peptides, in orange, are the selected Glu-C peptides, and in red are the selected tryptic peptides.

The first step in PRM setup is the selection of the proteotypic peptides. We performed an in-silico digestion of the H1 somatic subtypes with two proteolytic enzymes widely used in MS experiments, trypsin and endoproteinase Glu-C (Glu-C). We identified 53 unique peptides with less than 25 residues and a monoisotopic mass above 600 Da (Table S6). The higher number of unique peptides, 15, was found in H1X, the more divergent subtype. The lower number of candidates, four peptides, was obtained for H1.4, which shares a high sequence identity with H1.2 and H1.3.

Then, we analyzed the empirical suitability score (ESS) derived from the Peptide Atlas database, which represents a ranking of how suitable the peptide is as a reference or proteotypic peptide and the total number of observations in the current build of the database. Peptides with high ESS values (ESS > 0.85) belonged to H1.0, H1.5, and H1X. Peptides derived from the four remaining subtypes had ESS below 0.5. The ESS score includes penalties if the peptides have undesirable residues that impact a peptide’s suitability for targeting in PRM experiments, like the methionine residue in the highest-ranking peptide of H1.0 [32].

Following the in-silico analysis, we performed untargeted MS/MS experiments using a control sample containing all somatic H1 subtypes (Table S6). This sample was a mixture of perchloric acid extractions of three cell lines, HeLa, K652, and T47D, digested with the selected enzymes. We analyzed whether the peptide could be detected in the control sample, allowing us to discard approximately 20 from the initial 53 candidates, mainly corresponding to peptides with less than nine residues. We also checked the presence of the most frequent PTMs found in H1, phosphorylation, methylation, and acetylation. In most peptides no PTMs were detected, while those containing the N-terminal residue were always modified. Some miscleavaged peptides were detected, particularly with Glu-C, so the digestion conditions were optimized as described in material and methods. Some tryptic peptides of the RD– subtypes H1.1-H1.4 with miscleavages were selected for further analysis because they were the only species detected or their presence was much higher (≥90%) than the complete digestion product (Table S6). Considering these data, three peptides per subtype, except for H1.2 (two peptides), were synthesized containing heavy isotopes (Figure 1B; Table S7).

Several parameters were evaluated using the isotopically labeled peptides before selecting the quantification peptides: linearity between peptide concentration and peak area, MS/MS fragmentation pattern, and the shape of the extracted ion chromatogram (XIC) (Figures S1, S2, and Table S7). All tryptic peptides of H1.2-H1.4 were discarded because of the low r^2^ in the calibration curves and the broad shape (more than 20 minutes retention time, data not shown) of the peaks was not suitable for accurate quantification. Therefore, quantification of these subtypes was carried out using Glu-C peptides derived from the N-terminal domain, but not containing the N-terminal residues (Figure 1). Selected peptides for H1.2 and H1.4 have identical amino acid composition and mass, but they could be easily distinguished in PRM by their retention time and product ions (Figure S2). For the rest of the subtypes, the peptide with the highest ESS was selected for quantification (Figure 1B; Table S6). In the case of H1.0, the selected peptide contained a methionine residue, which is not optimal as a proteotypic peptide because it can be oxidized [31]. However, the rest of H1.0 peptides had miscleavages or low detectability, so we decided to select this peptide and correct the quantification considering both the oxidized and non-oxidized peptides.

We performed new calibration curves of the selected peptides with the control sample as a matrix. In these conditions, the calibration curves displayed a high degree of linearity with adjusted r^2^ > 0.98, and the limits of detection (LOD) and quantification (LOQ) were calculated (Table 2; Figure S3) [33]. During the PRM setup, we used perchloric acid extraction because it increased the amount of H1 peptides in the sample, so we verified that the proportions of the different subtypes were in agreement to those obtained using a total protein extract (Figure S4, Table S8).

**Table 2.** Peptides used in the Parallel Reaction Monitoring (PRM) assay development.

| Subtype | Number of candidates | Number of heavy peptides | Peptide sequence | r<sup>2</sup> calibration curve | LOD (ng/μg extract) | LOQ (ng/μg extract) |
| --- | --- | --- | --- | --- | --- | --- |
| H1.0 | 7 | 3 | YSDMIVAAIQAEK | 0.984 | 0.055 | 0.168 |
| H1.1 | 7 | 3 | KKPAGPSVSELIVQAASSSK | 0.993 | 0.037 | 0.111 |
| H1.2 | 6 | 2 | TAPAAPAAAPPAE | 0.996 | 0.026 | 0.08 |
| H1.3 | 6 | 3 | TAPLAPTIPAPAE | 0.998 | 0.02 | 0.06 |
| H1.4 | 4 | 3 | TAPAAPAAPAPAE | 0.995 | 0.032 | 0.098 |
| H1.5 | 8 | 3 | ATGPPVSELITK | 0.989 | 0.046 | 0.14 |
| H1X | 15 | 3 | ALVQNDTLLQVK | 0.996 | 0.027 | 0.066 |

LOD: Limit of detection
LOQ: Limit of quantification

### Quantification of H1 subtypes in human cell lines by PRM

To evaluate our PRM assay performance in the quantification of H1 complement, we used three human cell lines: HeLa, K562, and T47D, with different subtype compositions. Using the limits of quantification calculated with the pool of three cell lines, we quantified six subtypes in HeLa, five in K562, and six in T47D (Table S9, Figure 2A, Figure 3A). All subtypes, except H1.3, could be quantified in HeLa. H1.0 and H1X were present in low amounts, while H1.4 and H1.5 were the more abundant subtypes. In K562, five subtypes were quantified, H1.2-H1.5, and H1X. The most abundant subtype was H1.2, followed by H1.4. All subtypes, except H1.1, were quantified in T47D. In this cell line, the most abundant subtype was H1.5. Triplicates showed variation coefficients lower than 15%, indicating the reproducibility of the PRM measurements.

**Figure 2.**
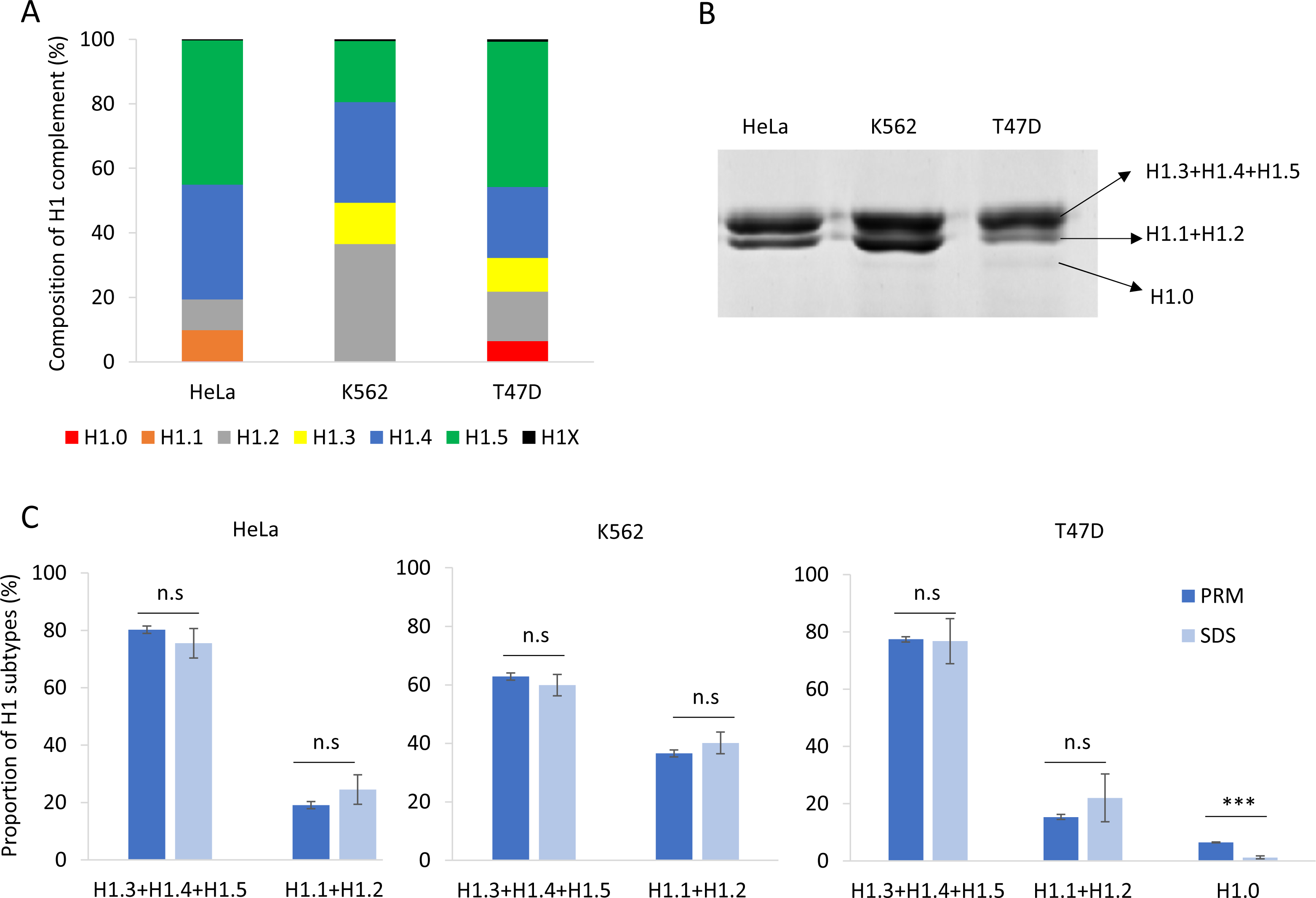
Quantification of the H1 complement in human cell lines. A. Composition of H1 complement in HeLa, K562, and T47D measured by PRM. Values correspond to the average of three replicates and are expressed as a percentage of the total H1 content. B. SDS-PAGE profile of perchloric acid extractions of HeLa, K562, and T47D. On the right is the expected subtype composition of each band. C. Quantification of the bands observed in perchloric acid extractions compared to the values obtained using PRM results. Error bars correspond to the standard deviation of three replicates. The differences between the PRM and SDS results were as described in materials and methods. n.s. not significat. *** Benjamini-Hochberg adjusted P-value< 0.001.

**Figure 3.**
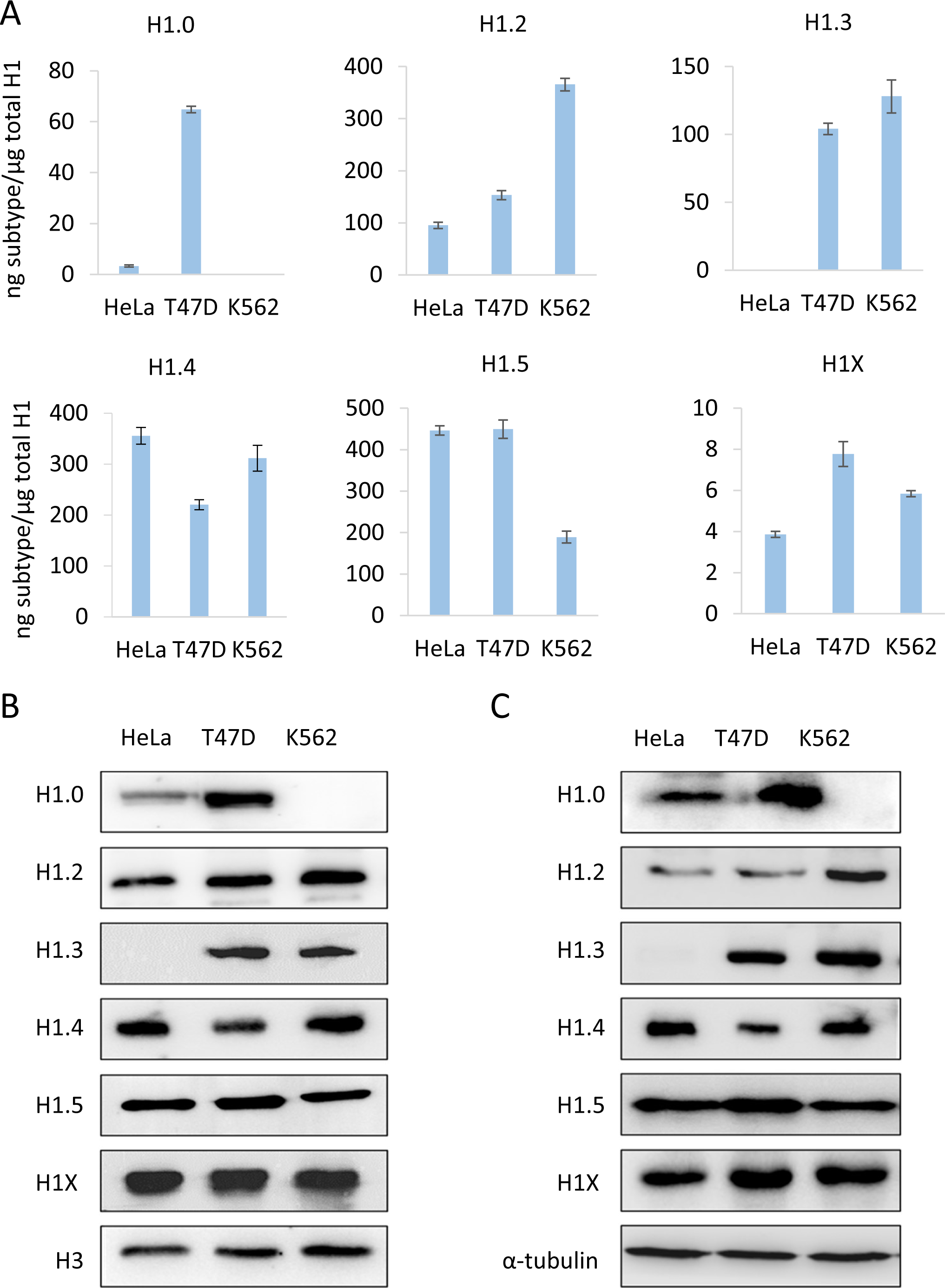
Abundance of individual H1 subtypes in three human cell lines, HeLa, T47D, and K562. A, quantification of H1 subtypes by PRM. B and C, analysis of H1 subtypes by western blot of sulfuric acid (B) and total protein (C) extracts. Error bars represent the standard deviation of the triplicates. Histone H3 and tubulin were used as loading controls for acid and total extracts, respectively.

Perchloric acid extraction followed by denaturing polyacrylamide gel electrophoresis allowed us to visualize the H1 complement separated into three bands (Figure 2B). The band with higher molecular weight contains three subtypes, H1.3-H15, while the band with intermediate molecular weight contains H1.2 and, when present, H1.1. The band with lower molecular weight, which is not always detectable, corresponds to H1.0. The subtype H1X is present in low proportions and must be detected by other techniques, such as Western blot. We used the proportions of H1 subtypes obtained by PRM to estimate the percentage of H1 expected in each band of the perchloric extraction for the three cell lines. Statistical comparison of the estimated proportions of H1 subtypes by PRM quantification with those obtained experimentally from SDS-PAGE resulted in no significant differences, except for H1.0. In the latter case, the discrepancy could be explained by the insufficient staining of low-abundance protein bands. As expected, the variability of the SDS-PAGE estimates was higher than that of PRM values (Figure 2C).

To further evaluate the PRM results, we compared the PRM data of the individual subtypes with Western blot from sulfuric and total protein extractions (Figure 3). The detection of H1 subtypes by Western blot is specific, however, the limit of detection depends on the antibody performance.

For instance, H1X was readily detected by Western blot despite being one of the less abundant subtypes. On the other hand, H1.1 is present in significant proportions in HeLa, but it was not detected by Western blot. Overall, Western blot results using both types of extracts agreed with the quantitative profile obtained by PRM, indicating the accuracy of the assay.

### Quantification of H1 subtypes from human samples

One of the main objectives of our study is to develop an absolute quantitative assay capable of evaluating the potential of H1 subtypes as biomarkers in disease using biological samples. As a proof of concept, we analyzed the H1 complement in the peripheral blood of healthy individuals and chronic myeloid leukemia patients. Due to the limited amount of protein samples available, the quantification was performed using one point of the calibration curve per sample. Peaks for quantification were manually validated. We scanned the MS1 spectra and detected the presence of miscleavaged peptides of several H1 subtypes, so we used an external calibration curve using the light synthetic miscleavaged peptides to correct quantification results (Table S10). External calibration curves were not acquired the same day as the samples.

We analyzed peripheral blood samples of healthy individuals with normal proportions of nucleated cell populations in white blood cells by PRM (Figure 4A). Quantification of H1 subtypes showed a similar amount of total H1 per mL of sample (Table S11). The composition of the H1 complement was also alike in the three individuals tested. The more abundant subtypes were H1.4 and H1.5, followed by H1.0 and H1.3. Subtypes H1.1, H1.2, and H1X were present in small amounts, representing between 1.5-5% of the total H1 (Figure 4B). However, the exact proportion of each subtype showed some variability among individuals.

**Figure 4.**
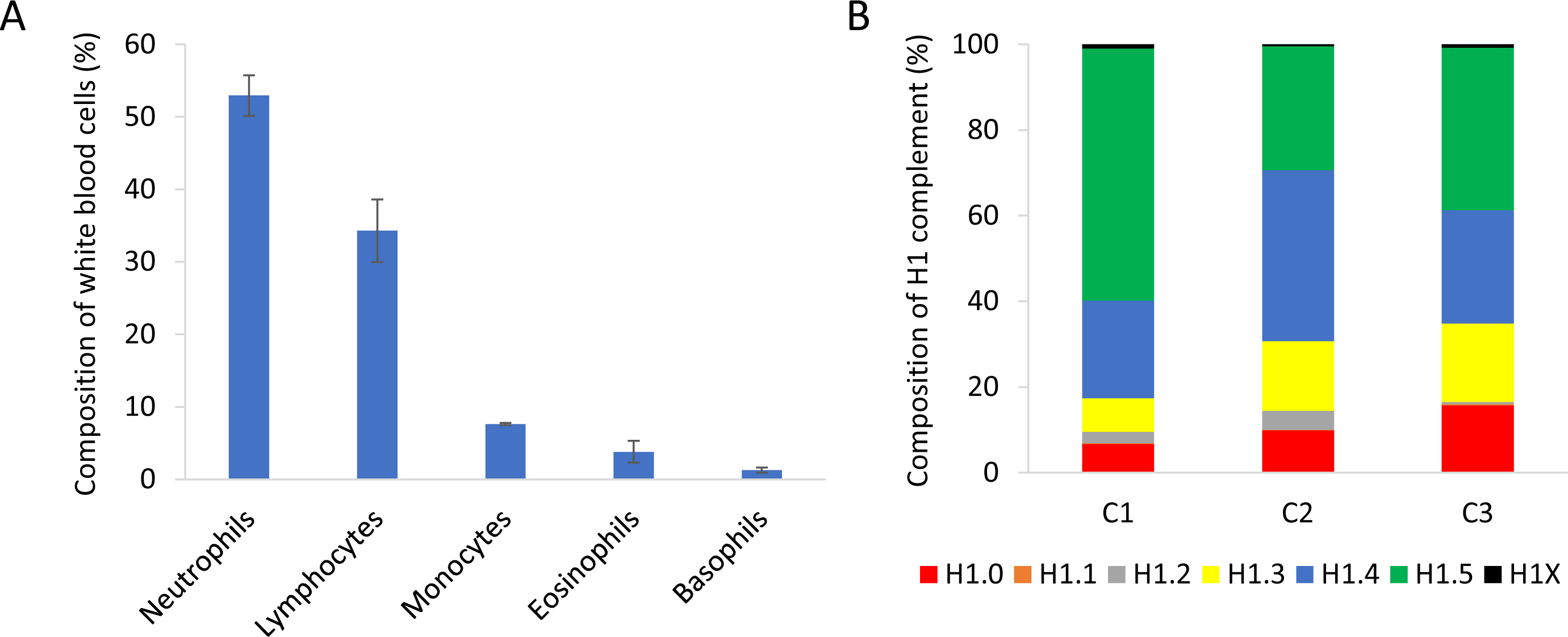
Analysis of peripheral blood samples of healthy individuals. A. Percentages of the different nucleated cell populations present in peripheral blood. The values showed the average of the three healthy individuals included in the study. Error bars correspond to the standard deviation. B. Composition of the H1 complement quantified by PRM in each healthy control sample (C1, C2, and C3).

We also quantified H1 subtypes from peripheral blood cells of eight patients diagnosed with chronic myeloid leukemia. The age of the patients ranged from 35-75 years, and most of them belonged to the low-risk category. The small cohort included six patients who responded to therapy with imatinib and two who didn’t (Table S12). We addressed two objectives. First, to study if one or several H1 subtypes could contribute to the resistance to TKI. Second, to explore whether H1 subtypes could aid in predicting the response to tyrosine kinase inhibitors (TKIs).

The complement of H1 showed high variability among patients (Figure 5A, Table S13). In six patients, only one subtype, H1.0 in two and H1.5 in four, accounted for more than 50% of the total H1. Meanwhile, the other two patients had a more heterogeneous H1 complement. Subtype H1.2 was quantified in only one patient. We analyzed if the non-responder patients had similarities in the proportions of some H1 subtypes. We found by clustering analysis that subtypes H1.0, H1.1, and H1X grouped non-responder patients in a tight cluster, characterized by lower proportions of these three subtypes than most responders to TKIs (Figure 5B). Principal component analysis confirmed the clustering results, grouping non-responders together (Figure 5C). These results suggest that one or more H1 subtypes may be involved in imatinib resistance.

**Figure 5.**
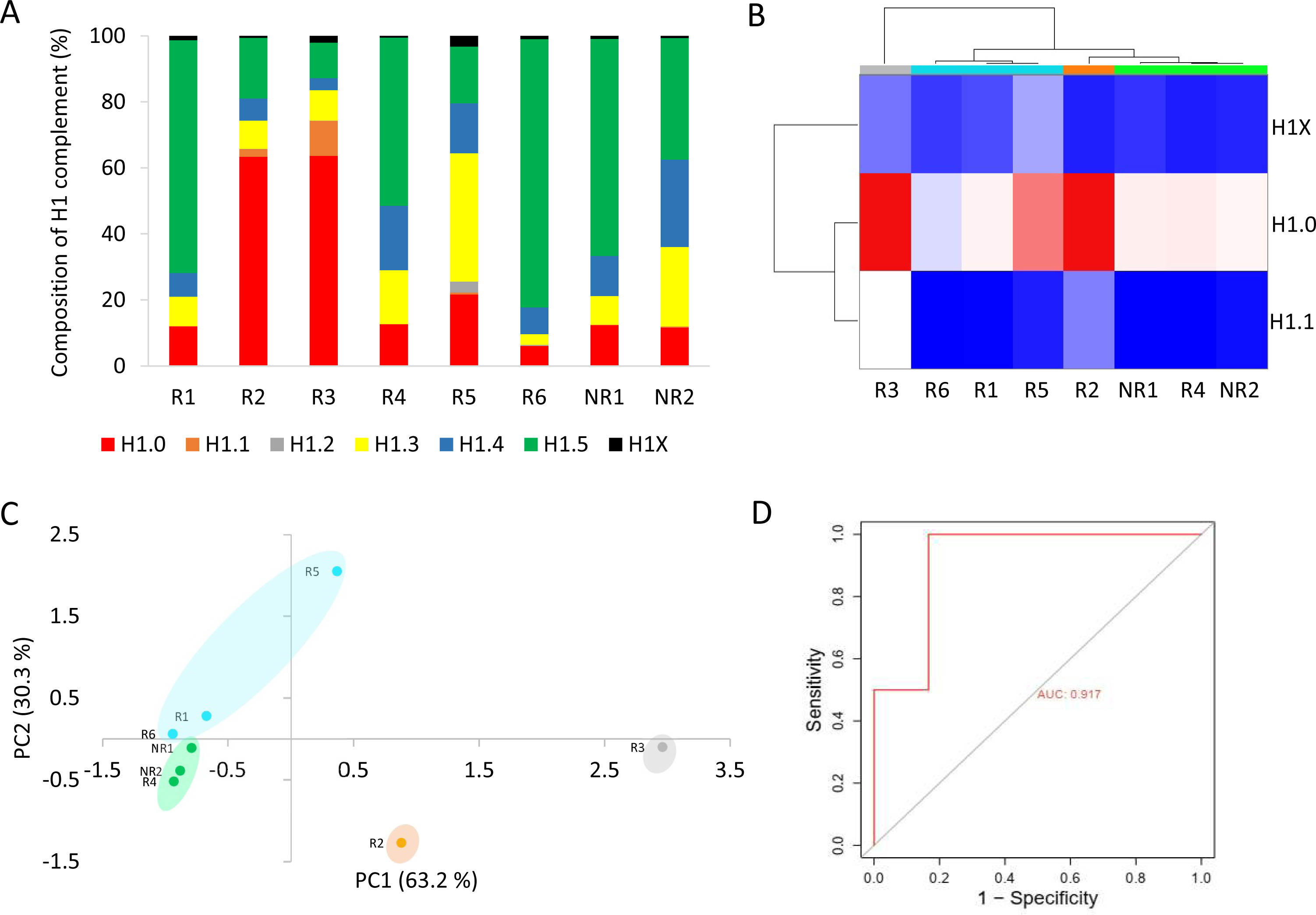
Analysis of H1 complement in CML patients by PRM. A. Composition of the H1 complement quantified by PRM in CML patients. B. Heatmap of the proportions of H1.0, H1.1, and H1X in CML patients obtained by hierarchical clustering based on correlation distances with average linkage. C. Principal component analysis (PCA) of the data used for hierarchical clustering. The colors correspond to those assigned to the different clusters in panel B. In parenthesis, the percentage of variation is explained by each component. D. ROC curve evaluating the accuracy of the total H1 levels predicting imatinib response. R1-R6, responders to imatinib. NR1 and NR2, non-responder patients. AUC, area under the curve.

The total H1 per mL was also variable among CML patients (Table S13). However, we observed that the patients who did not respond to imatinib had higher values than most responders. To analyze the performance of a diagnostic test with high statistical reliability the cohort must be large and balanced [34]. Our cohort of CML patients did not meet these criteria. Nevertheless, as a prospective simulation, we analyzed the specificity, sensitivity, and accuracy of the total H1 quantification to predict imatinib response. We found that the total H1 levels predicted the response to therapy with a specificity of 0.833 and a sensitivity of 1, according to Youden’s index (J). The accuracy was evaluated using a receiver operating characteristic (ROC) curve, obtaining an area under the curve (AUC) of 0.915, which confirmed that the measurement of H1 levels could predict imatinib response (Figure 5D). These results, albeit preliminary, encourage further research and their validation in an independent and larger cohort.

## Discussion

Somatic cells can express up to seven different H1 subtypes, whose relative proportions compose the H1 complement. Subtype composition can vary in physiological conditions depending on several factors [4]. Changes in the proportions of most H1 subtypes have also been associated with disease, in particular with cancer[6–20]. Therefore, studying subtype composition may help to understand the functional role of individual H1s and their contribution to disease development. We have developed an MS-based parallel reaction monitoring assay using proteotypic peptides from all somatic H1 subtypes, suitable to characterize H1 complement in physiological and pathological conditions. In PRM, several proteins can be quantified together, decreasing experimental error. This approach has been used successfully for the quantification of H2A and H2B variants[35]. The high sequence identity between H1 subtypes made it necessary to use two different proteolytic enzymes to generate unique peptides for each subtype. This procedure could result in biased quantification due to the specific proteolysis efficiency of the two enzymes.

We evaluated the PRM assay by quantifying the H1 complement in three human cell lines. The results were reproducible among replicates, with variation coefficients below 15%, and they agreed with the values estimated from SDS-PAGE data, considering the expected composition of each band. The correspondence between SDS-PAGE and PRM results point out that using two enzymes had little or no effect on the quantification results. Moreover, the relative amounts in the three cell lines of individual subtypes analyzed by Western blot confirmed PRM results. These results suggest that our PRM assay is accurate and reproducible. In addition, PRM is highly sensitive and needs low protein amounts, which makes it ideal for analyzing H1 in biological samples. The calculated limits of quantification (LOQ) for acid extracts of human cell lines were below 0.17 ng, supporting this conclusion.

As a proof of concept, we characterized the H1 complement of circulating white blood cells of healthy human subjects and CML patients. In healthy patients, the amount of H1 per mL of sample was relatively similar, and the more abundant subtypes were H1.4 and H1.5. Individual differences were observed in the H1 complement, which could be explained, at least in part, by the different cell type proportions in white blood cells.

We analyzed a small cohort of CML patients, with 20% of imatinib-resistant patients. This percentage is consistent with the one reported for this disease[25,26]. We found that those patients had a higher content of H1 per mL of sample. This parameter predicted the response to imatinib with 83.3% specificity and an AUC of 0.917 in the ROC curve. These results could be misleading and overly optimistic due to the limited number of subjects and the presence of more negative (responders to imatinib) samples than positive ones (non-responders)[34]. However, the increase in the global content of H1 has already been associated with the tumor proliferation rate in prostate cancer, while in glioblastoma, lower levels of H1 correlated with low survival rates[9,11].

Resistance to TKIs in CML is associated with different phenomena, including mutations in the fusion protein BCR-ABL and several chromatin-related processes in which H1 may be involved, such as DNA repair, genome instability, and epigenetic dysfunction [25,36–39]. The composition of the H1 complement could be relevant for resistance development, so we analyzed the similarities in the proportions of specific subtypes between non-responders. Clustering and PCA analysis showed that the patients who didn’t respond to imatinib, despite having higher H1 content, had lower relative proportions of H1.0, H1.1, and H1X.

Previous studies have associated changes in H1.0, H1.1, and H1X with cancer progression. In the case of H1.0, low levels have been associated with high proliferative activity and a poor outcome in breast cancer [7,8]. A low expression of H1.0 has been observed in cells with stem-like properties within tumors. These cells were characterized by long-term proliferation and metastatic potential[36]. On the other hand, in ovarian cancer, H1.0 upregulation is associated with therapy resistance, disease recurrence, and poor survival [20]. Lower levels of H1.1 were present in prostate adenocarcinomas when compared with healthy tissues. The role of this subtype in prostate tumorigenesis was associated with the modulation of the Wnt signaling pathway [13].

Finally, increased expression of H1X has been described as a favorable prognosis biomarker in astrocytic gliomas, while it was also associated with neuroendocrine tumors [6,10].

Several studies link H1 subtypes to hematopoiesis and hematological malignancies, supporting the possibility that they may contribute to imatinib resistance. Transcription factors involved in hematopoiesis bind to the promoter of H1 genes, probably controlling their expression, whereas the knockdown of different H1 subtypes altered neutrophil differentiation [40,41]. Moreover, decreased expression of H1.3 is associated with a bad prognosis in acute myeloid leukemia patients with mutations in NPM1, while mutations in several RD subtypes are recurrent in B-cell lymphomas [16,42]. Our preliminary results hint that the levels of H1 may contribute to predicting imatinib response, while several H1 subtypes may be associated with the resistance to this drug. Further studies are needed to corroborate our findings in a larger cohort and to unveil the underlying molecular mechanisms in the role of H1 in imatinib resistance.

## Conclusions

We have designed a PRM assay for quantifying human H1 subtypes, suitable for characterizing the H1 complement in physiological conditions and to evaluate their potential as cancer biomarkers. Our PRM assay was validated using three cancer cell lines showing high sensitivity, reproducibility, and specificity. We have used our PRM assay to characterize the H1 complement of circulating white blood cells in healthy subjects and CML patients. As proof of concept, we analyzed the association between the H1 content and imatinib response. Our results suggest that the levels of H1 might predict imatinib response. However, these preliminary findings require confirmation in a large and independent cohort to test the suitability of this parameter as a biomarker for therapy response in CML. We also found that the H1 complement of non-responders is characterized by lower proportions of H1.0, H1.1, and H1X, suggesting that the absence of these subtypes may contribute to the development of TKI resistance.

## Declarations

### Ethics approval and consent to participate

Human samples of peripheral blood from three healthy individuals and eight patients of CML were obtained following the ethical guidelines of the Germans Trias i Pujol Hospital.

### Consent for publication

Not applicable.

### Availability of data and materials

All data generated or analyzed during this study are included in this published article, its supplementary information files, and in the ProteomeXchange Consortium via the PRIDE partner repository with the dataset identifier PXD050243.

## Authors’ contributions

LGJ prepared the protein extracts, performed the experiments for the validation of the PRM, and contributed to the analysis of the data. VL performed the mass spectrometry experiments, standardized the PRM assay, carried out the quantification, and contributed to writing the manuscript. AM prepared protein extracts of human cell lines and peripheral blood samples. PI supervised the sample preparation, the validation of the PRM assay, and contributed to the conceptualization of the study and in writing the manuscript. XB, provided the clinical perspective and data on CML. ZL, provided and processed the peripheral blood samples, and contributed to the conceptualization of the proof of concept using CML patients. VM supervised all the proteomics experiments and data analysis, contributed to the conceptualization of the study and in writing the manuscript. AR supervised the analysis of the data, was a major contributor to the conceptualization of the study and in writing the manuscript and secured the funding to carry out the study. All authors read and approved the final manuscript.

## Abbreviations

ACN: Acetonitrile
AUC: Area under the curve
CML: Chronic myeloid leukemia
DTT: Dithiothreitol
EDTA: 2,2’,2’’,2’’’-(ethane-1,2-diyldinitrilo) tetraacetic acid
emPAI: exponentially modified abundance index
ESS: Empirical suitability score
FA: Formic acid
FDR: False discovery rate
Glu-C: Endoproteinase
Glu-C IAM: Iodoacetamide
iBAQ: intensity based absolute-quantitation
LH: Light/Heavy
LOD: Limit of detection
LOQ: Limit of quantification
MS: Mass spectrometry
PBS: Phosphate-buffered saline
PCA: Principal component analysis
PIC: Protease inhibitor cocktail
PRM: Parallel reaction monitoring
PVDF: Polyvinylidene difluoride
RD: Replication-dependent
RI: Replication-independent
ROC: Receiver operating curve
SDS: Sodium dodecyl sulfate
SDS-PAGE: Denaturing polyacrylamide gel electrophoresis
TKI: Tyrosine-kinase inhibitor
XIC: Extracted ion chromatogram

## Acknowledgements

We thank Marina Gay from the Institute for Research in Biomedicine (IRB Barcelona) for her help with the analysis of the PRM data.

