## Supplementary Figures for "Quantification of histone H1 subtypes using targeted proteomics"

Figure S1

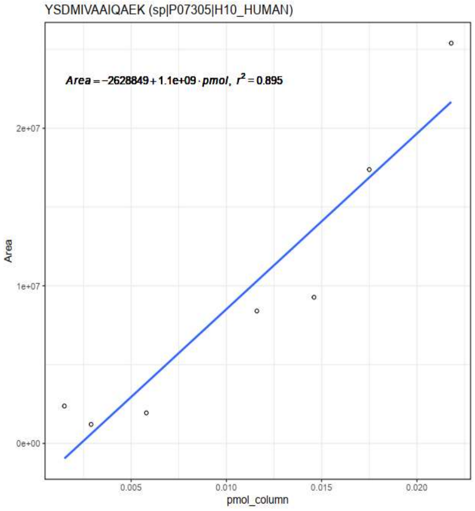


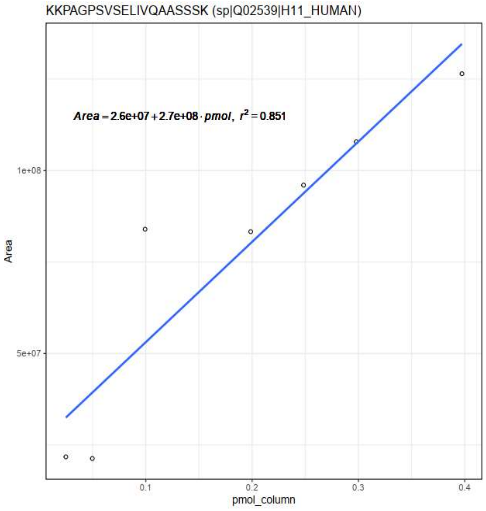


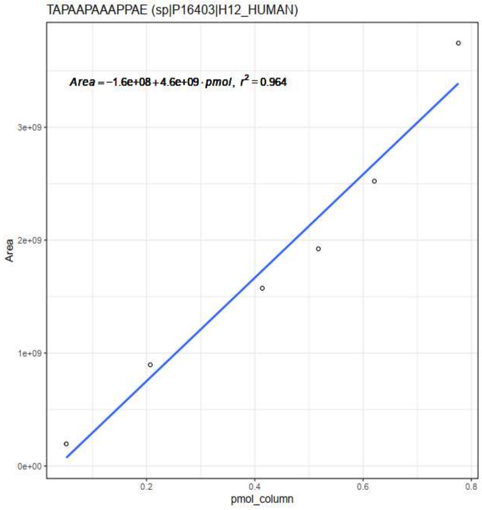


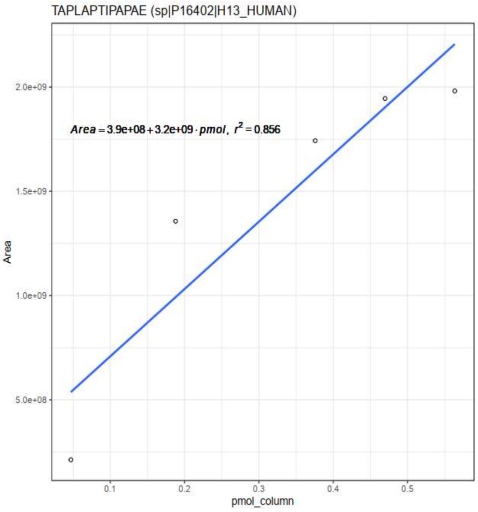


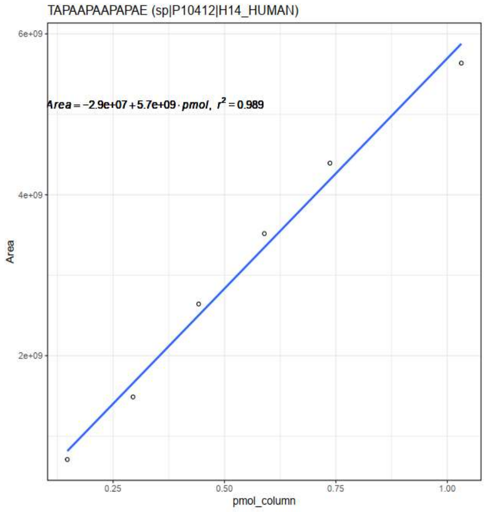


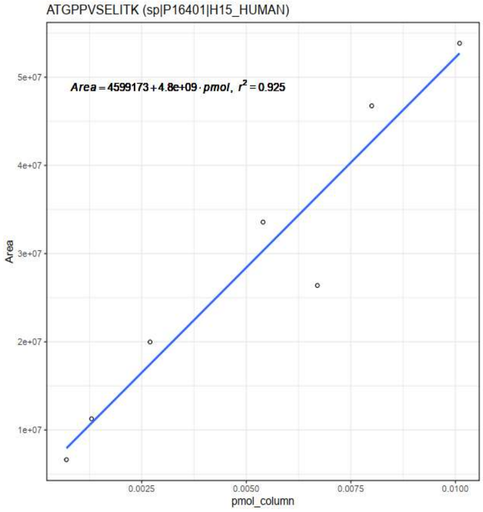


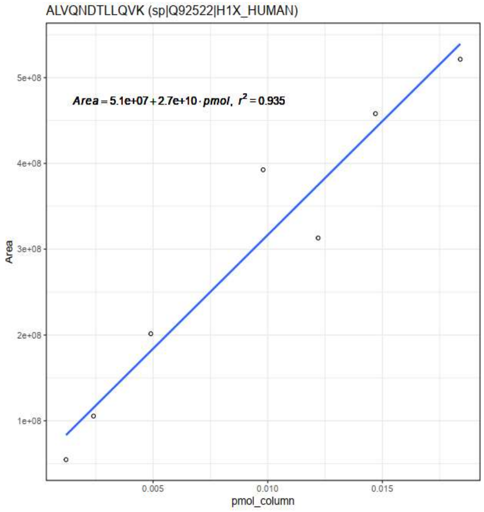


**Figure S1. Initial calibration curves of the heavy peptides selected for quantification of H1 subtypes.** Peptides were disolved in 3% ACN and 1% FA, as described in materials and methods.

Figure S2

A

H1.2: TAPAAPAAAPPAE; m/z 567.7931

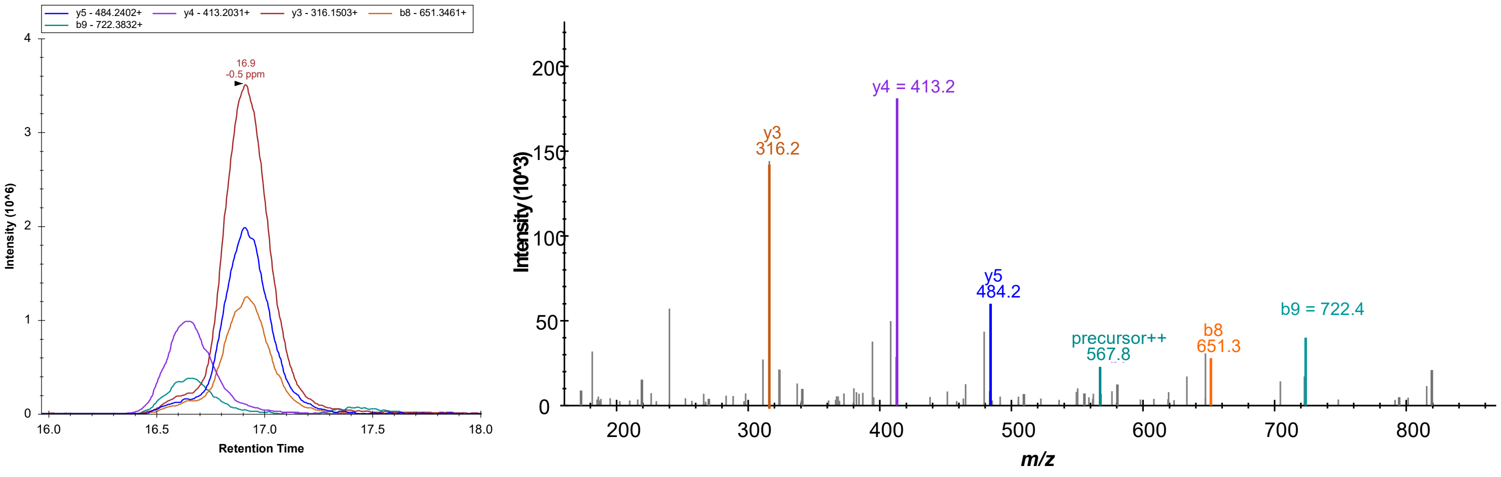


B

H1.4: TAPAAPAAPAPAE; m/z 567.7931

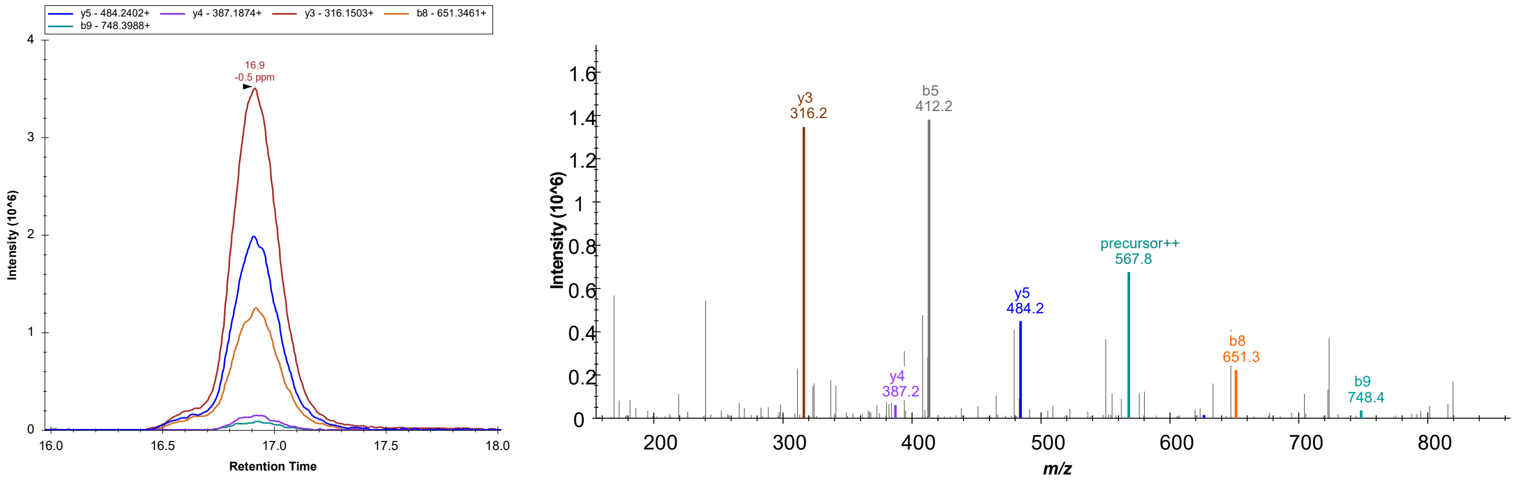


**Figure S2. Analysis of H1.2 and H1.4 selected peptides.** Extracted ion chromatogram of different transitions from Skyline and fragmentation spectra of, A. H1.2 selected peptide and B. H1.4 selected peptide. Only transitions y4 and b9 were used to quantify these two peptides, as they are the only transitions showing different masses.

Figure S3

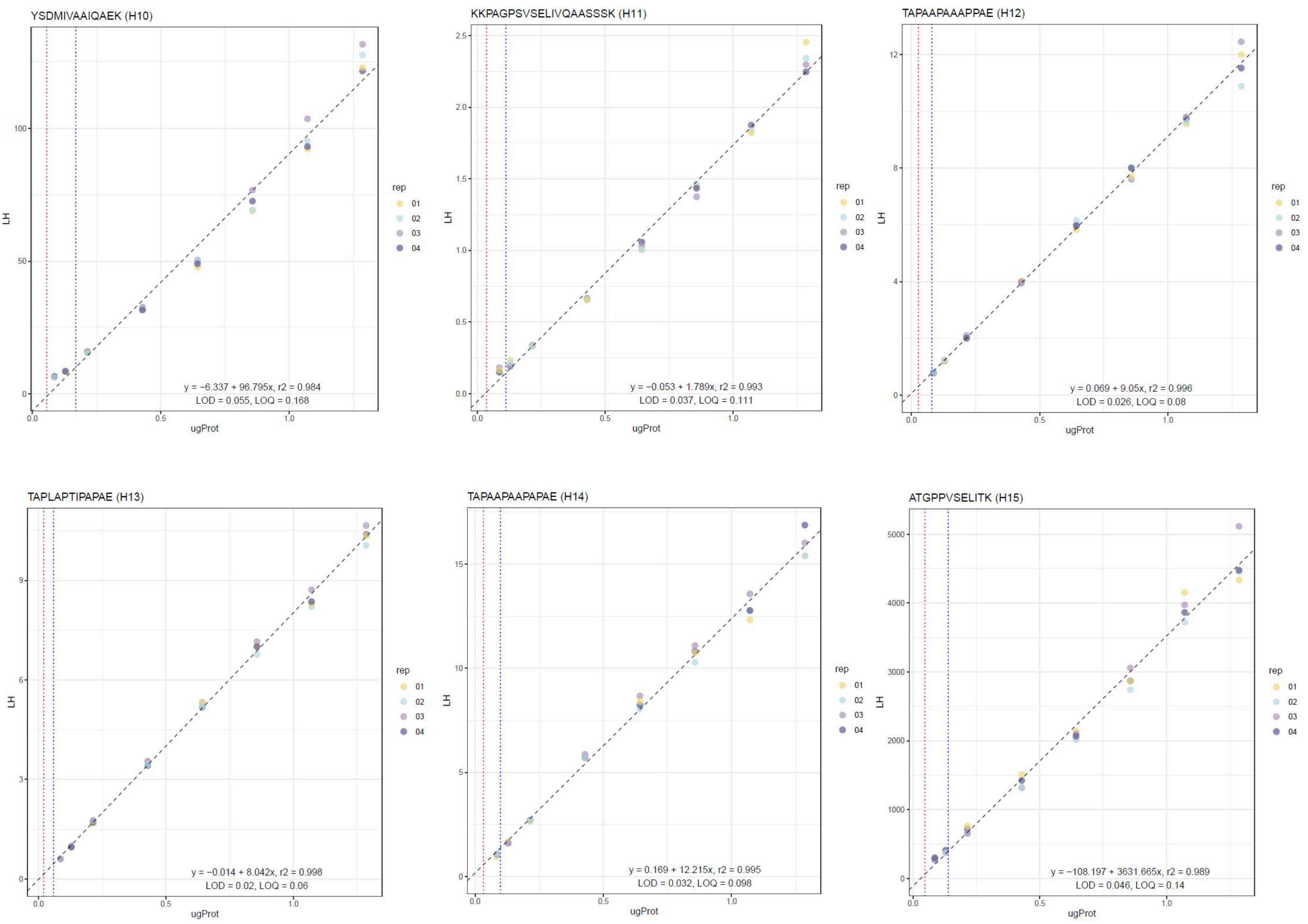


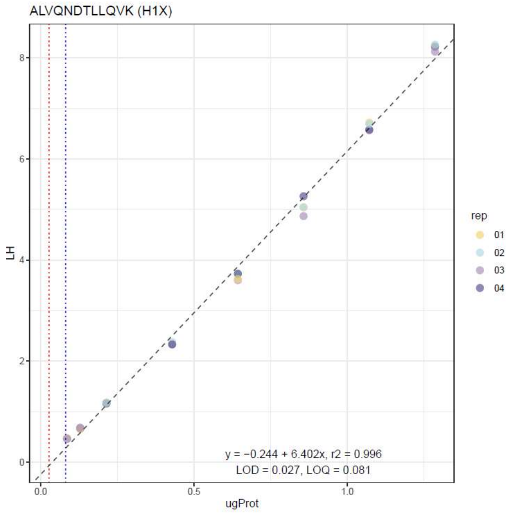


**Figure S3. Calibration curves of the heavy peptides used for quantification of H1 subtypes.** All the curves were obtained using four replicates of each data point. The equation shown was used to calculate the limits of detection (LOD) and quantification (LOQ), as described in materials and methods.

Figure S4

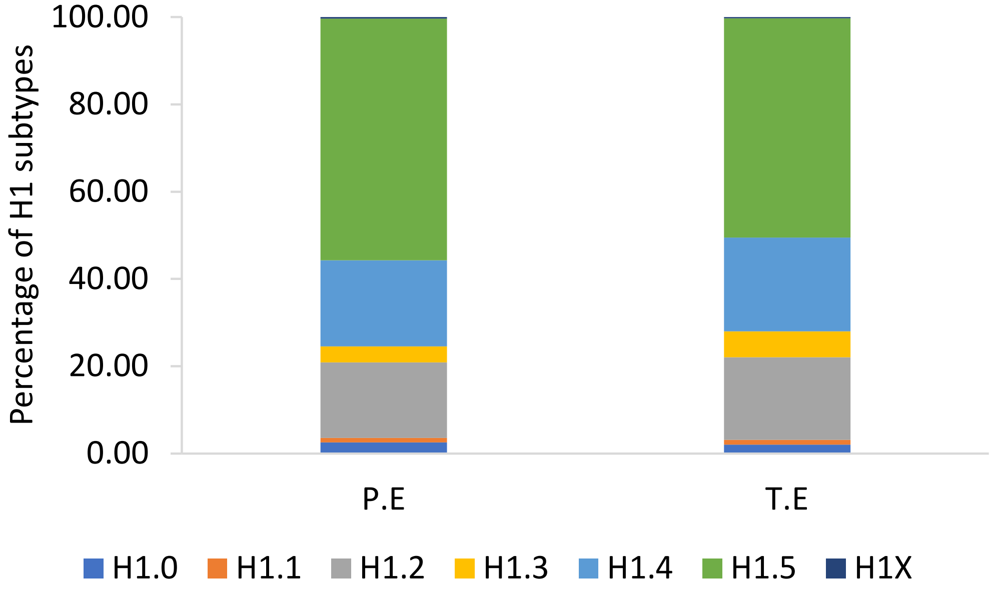


**Figure S4. Proportions of H1 subtypes in the pool sample quantified from perchloric acid extracts (P.E) and total protein extracts (T.E).**
